# Efficient Neutralization of SARS-CoV-2 Omicron and Other VOCs by a Broad Spectrum Antibody 8G3

**DOI:** 10.1101/2022.02.25.482049

**Authors:** Hang Ma, Chien-Te K. Tseng, Huifang Zong, Yunji Liao, Yong Ke, Haoneng Tang, Lei Wang, Zhenyu Wang, Yang He, Yunsong Chang, Shusheng Wang, Aleksandra Drelich, Jason Hsu, Vivian Tat, Yunsheng Yuan, Mingyuan Wu, Junjun Liu, Yali Yue, Wenbo Xu, Xiaoju Zhang, Ziqi Wang, Li Yang, Hua Chen, Yanlin Bian, Baohong Zhang, Haiyang Yin, Yi Chen, En Zhang, Xiaoxiao Zhang, John Gilly, Tao Sun, Lei Han, Yueqing Xie, Hua Jiang, Jianwei Zhu

## Abstract

Numerous mutations in the spike protein of SARS-CoV-2 B.1.1.529 Omicron variant pose a crisis for antibody-based immunotherapies. The efficacy of emergency use authorized (EUA) antibodies that developed in early SARS-CoV-2 pandemic seems to be in flounder. We tested the Omicron neutralization efficacy of an early B cell antibody repertoire as well as several EUA antibodies in pseudovirus and authentic virus systems. More than half of the antibodies in the repertoire that showed good activity against WA1/2020 previously had completely lost neutralizing activity against Omicron, while antibody 8G3 displayed non-regressive activity. EUA antibodies Etesevimab, Casirivimab, Imdevimab and Bamlanivimab were entirely desensitized by Omicron. Only Sotrovimab targeting the non-ACE2 overlap epitope showed a dramatic decrease activity. Antibody 8G3 efficiently neutralized Omicron in pseudovirus and authentic virus systems. The in vivo results showed that Omicron virus was less virulent than the WA1/2020 strain, but still caused deterioration of health and even death in mice. Treatment with 8G3 quickly cleared virus load of mice. Antibody 8G3 also showed excellent activity against other variants of concern (VOCs), especially more efficient against authentic Delta plus virus. Collectively, our results suggest that neutralizing antibodies with breadth remains broad neutralizing activity in tackling SARS-CoV-2 infection despite the universal evasion from EUA antibodies by Omicron variant.

## Introduction

As a single-stranded RNA virus, the mutagenic nature of severe acute respiratory syndrome coronavirus 2 (SARS-CoV-2) has well been recognized since the initial global pandemic of coronavirus disease 2019 (COVID-19). The mutation rate of SARS-CoV-2 is ∼10^−6^ mutations/site/cycle, which is sufficient to cause more than one mutation per site during an infection period^1^. Especially, the frequency of manifold mutations in the spike (S) region is three times that of the entire genome^2^. With the vaccination starting from the end of 2020, the pandemic has been effectively alleviated. However, vaccines and therapeutic antibodies designed and developed based on the early circulating strains are inadequate to deal with the diversified virus mutations, especially the emergence of variants of concern (VOCs). Recently, the B.1.1.529 variant labeled with Omicron as the fifth VOC strain by world health organization (WHO) has caused great concern. Universal mutation sites were found in the S protein of Omicron and 15 of which are in the receptor binding domain (RBD). Those mutations reshapes the spatial structure of S protein more severely than any other VOC strain before^3^. After analyzing the cryo-EM structure of the trimeric S protein of the Omicron variant, researchers found that these mutations increased the interaction between the adjacent “up” conformation RBD and the “down” conformation RBD, making the S conformation less heterogeneous^4^. Besides, the affinity of Omicron spike protein binds to the ACE2 receptor is nearly 10 times higher than that of the wild type^5^. Cao et al. investigated the activity of nine emergency use authorized (EUA) antibodies against Omicron strain and found that most of them lost neutralization, and only Sotrovimab (VIR-7831) and DXP-604 retained moderate activity with pseudovirus neutralization IC_50_s of 0.181 and 0.287 μg/mL, respectively^6^. Another study also came to similar conclusions, only Brii-198 showed an enhanced neutralization with IC_50_ = 0.1-1 μg/mL against Omicron^7^.

In this study, we used a neutralizing antibody repertoire that was developed from COVID-19 convalescents at early 2020 and panned potent antibodies against Omicron. One of which, antibody 8G3 exhibited excellent Omicron neutralizing activity, as well as showed broad-spectrum against other variants in both pseudo- and authentic-virus systems. Currently, few studies have compared the activity of EUA antibodies against Omicron variant in the authentic virus system. We performed micro-well authentic virus neutralization and found that most EUA antibodies lost their activity against Omicron, only Sotrovimab retained partial neutralization. The newly found antibody 8G3 showed potent neutralizing activity and was 47 times more efficient than Sotrovimab against Omicron. Thus, although the emergence of Omicron variant brings tough challenge to the existing EUA antibodies, antibodies with breadth can still be developed as a solution.

## Results

### Screening of neutralizing antibodies against Omicron variant

Using an B cell antibody repertoire from early stage COVID-19 convalescents^8^, we compared dozens of candidate molecules in neutralizing activity against pseudo-typed SARS-CoV-2 WA1/2020 and Omicron viruses. We found that most of the antibodies that performed well against the WA1/2020 virus nearly lost their neutralization against Omicron; only a few antibodies retained full or partial activity (Fig. 1a-b). Antibodies 9A10, 8C12, 13A12, 3A4, 2B8, 11D3, 9C4, 9B10, and 9E12 reduced the neutralization from 100% against WA1/2020 to less than 20% against Omicron at the concentration of 1 μg/mL. Only several antibodies, including 7G10, 8G4 and 9D11 retained 53%, 63% and 77% neutralization against Omicron at the concentration of 1 μg/mL, respectively. The most noteworthy is the antibody 8G3, which still achieved 100% neutralization against Omicron at this concentration. Subsequently, we explored the concentration-dependent neutralization using 8G3, 8G4 and 9D11. The neutralizing ability of 8G4 and 9D11 was both moderate, with IC_50_ values of 1.0670 and 0.6584 μg/mL, respectively (Fig. 1c). Antibody 8G3 showed a strong activity with as an IC_50_ value of 0.0117 μg/mL against Omicron. In addition, antibody 8G3 also showed good neutralization against other SARS-CoV-2 strains, with IC_50_ values of 0.0140, 0.0825, 0.0210, 0.0044, 0.0147 and 0.0219 μg/mL for WA1/2020, Alpha, Beta, Gamma, Delta and Kappa pseudoviruses, respectively (Fig. 1d).

**Fig. 1.**
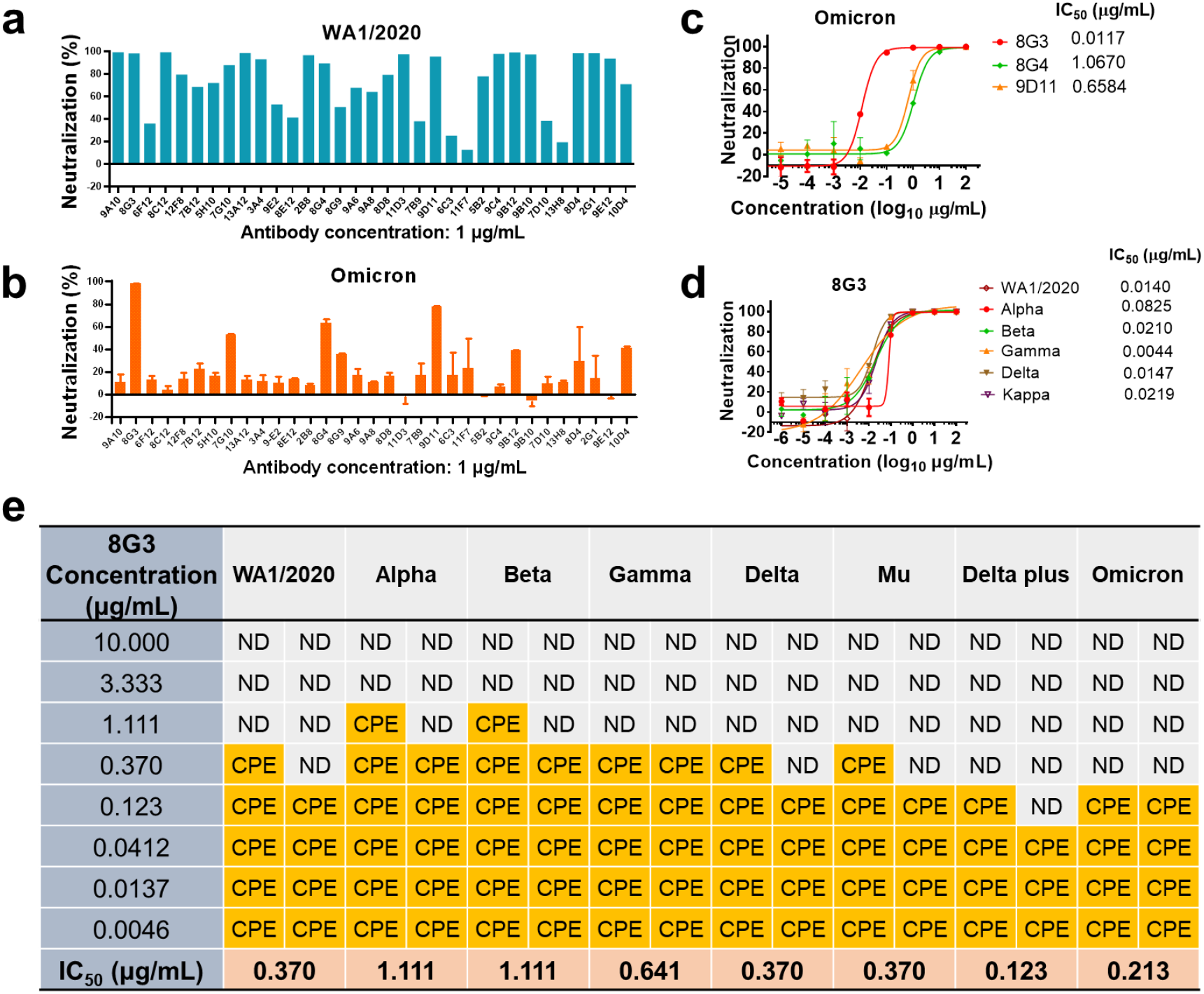
Screening of anti-SARS-CoV-2 Omicron antibodies from B cell antibody repertoire. **a-b**, Neutralization of WA1/2020 (a) and Omicron (b) pseudoviruses by antibody candidates. Pseudoviruses with active titer over 1×10^7^ TU/mL were mixed with antibodies at the final concentration of 1 μg/mL and used to infect the 293T-ACE2 cells. Two days later, the neutralizing efficacy was quantified according to the fluorescence intensity from luciferase-substrate reaction. **c-d**, Concentration-dependent neutralization against pseudo-typed SARS-CoV-2 Omicron (c) and other variants (d) by antibody candidates. Data are displayed as mean ± S.D. **e**, Dose-dependent authentic SARS-CoV-2 neutralization by antibody 8G3.

We then evaluated the authentic virus neutralizing activity of 8G3. The results showed that 8G3 neutralized WA1/2020, Alpha, Beta, Gamma, Delta, Mu, Delta plus, and Omicron viruses with IC_50_ values of 0.370, 1.111, 1.111, 0. 641, 0.370, 0.370, 0.123 and 0.213 μg/mL, respectively (Fig. 1e). Antibody 8G3 appeared to have higher activity against Delta Plus and Omicron compared with WA1/2020, which is a favorable sign for addressing current SARS-CoV-2 variants. Delta Plus has an additional K417N mutation in the RBD, but 8G3 was more efficient against Delta Plus than Delta. Thus the K417N is not a variate affecting 8G3 activity. These results suggested that antibody 8G3 has broad neutralizing effects against prevalent variants, and is possibly a conserved epitope antibody.

### Comparison of 8G3 and EUA antibodies

Subsequently, we compared antibody 8G3 with EUA antibodies in authentic wild type and Omicron virus neutralization. Antibody 8G3 showed potent activity against both WA1/2020 and Omicron viruses, with IC_50_ values of 0.080 and 0.137 μg/mL, respectively (Fig. 2a-b). EUA antibodies Etesevimab, Casirivimab, Imdevimab and Bamlanivimab neutralized WA1/2020 with high efficiency, but completely lost their activity against Omicron (IC_50_ > 100 μg/mL). Although the WA1/2020 neutralization ability of Sotrovimab was slightly inferior to other EUA antibodies, it retained partial activity against Omicron with an IC_50_ value of 6.409 μg/mL (Fig. 2b). By comparing the IC_50_s, Antibody 8G3 was about 47 times more potent than Sotrovimab.

**Fig. 2.**
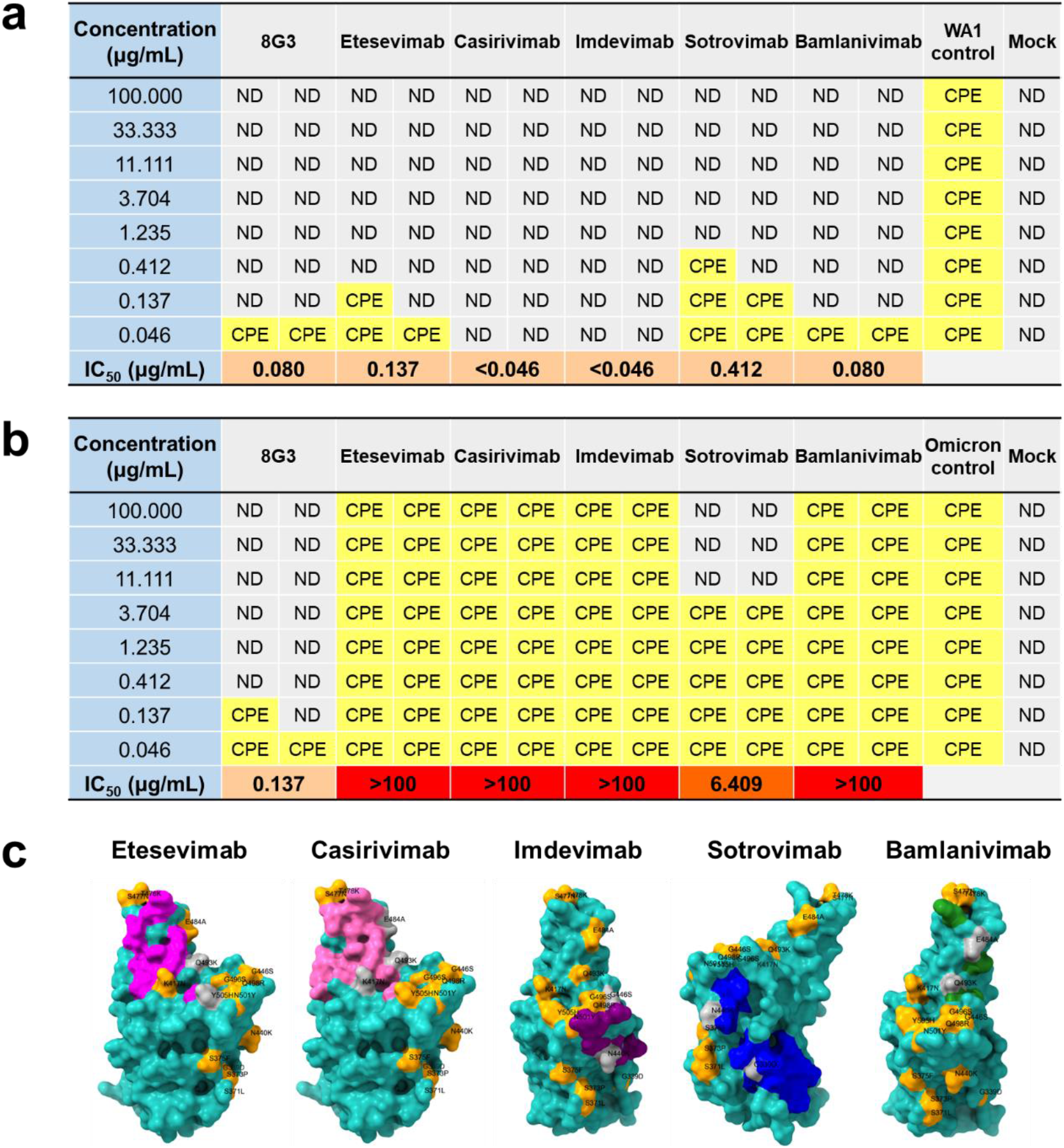
Authentic SARS-CoV-2 virus neutralization. **a-b**, Micro-well neutralization against authentic SARS-CoV-2 WA1/2020 (a) and Omicron (b) viruses with 3-fold serial diluted antibodies. **c**, The epitope surfaces of Etesevimab, Casirivimab, Imdevimab, Sotrovimab and Bamlanivimab on RBD are in magenta, pink, purple, blue and green, respectively. The mutation sites of Omicron are colored yellow, and are marked in dark gray if they overlap with antibody epitopes. CPE, cytopathic effect. ND, no detection.

The Omicron variant have 15 mutation sites widely distributed on the surface of the RBD. Combined with the authentic virus neutralization results, we can infer that antibodies targeting the RBD-ACE2 interface may be more susceptible to Omicron variant (Fig. 2c). Antibodies Etesevimab, Casirivimab and Bamlanivimab^9-11^, with epitopes highly overlapping with the RBD-ACE2 interface, completely lost their activities against Omicron. Sotrovimab, away from the RBD-ACE2 interface^12^, retained partial activity, while the G339D and N440K mutations within the epitope significantly reduced the neutralization by nearly 50 times (Fig. 2a-b). The epitope of antibody Imdevimab is at the edge of the RBD-ACE2 interface^10^, completely lost its activity possibly due to its small antigen contact area with N440K and G446S mutations.

### In vivo therapeutic efficacy

The protective efficacies of antibody 8G3 against the infection SARS-CoV-2 WA1/2020 and Omicron were evaluated in a human ACE2 transgenic mice model. Animals were challenged with 300 half tissue culture infectious dose (TCID_50_) of WA1/2020 virus or 2.3 × 10^4^ TCID50 of Omicron virus, and then treated with 20 mg/kg or 6.7 mg/kg of antibody 8G3 at 4h and 2 days post infection. Mice challenged with WA1/2020 virus and treated with PBS rapidly developed acute wasting syndrome and reached the endpoint on day 5. In contrast, both 20 mg/kg and 6.7 mg/kg of 8G3 effectively prevented weight loss, improved clinical scores, and completely cleared viral loads in lung and brain tissues of mice (Fig. 3a-c), which showed the good therapeutic ability of 2G1 against WA1/2020 virus. As for the Omicron virus infected mice, less body weight loss was observed although mice were challenged with a high viral load (Fig. 3a). There was fluctuations in clinical scores but only 1 mouse reached the designed endpoint (Fig. 3b). In addition, the viral load in the lungs of the control group on day 4 was lower than that of mice infected with WA1/2020, and no virus was detected in the brain. The virus was undetectable both in the lungs and brain on day 7 (Fig. 3c). These results suggested that the virulence of Omicron may reduced comparing with the WA1/2020 strain. Treatment of Omicron-infected mice with antibody 8G3 prevented adverse clinical manifestations, and completely cleared viral load in the lungs, suggesting a good therapeutic ability of 8G3 to protect animals from Omicron virus infection.

**Fig. 3.**
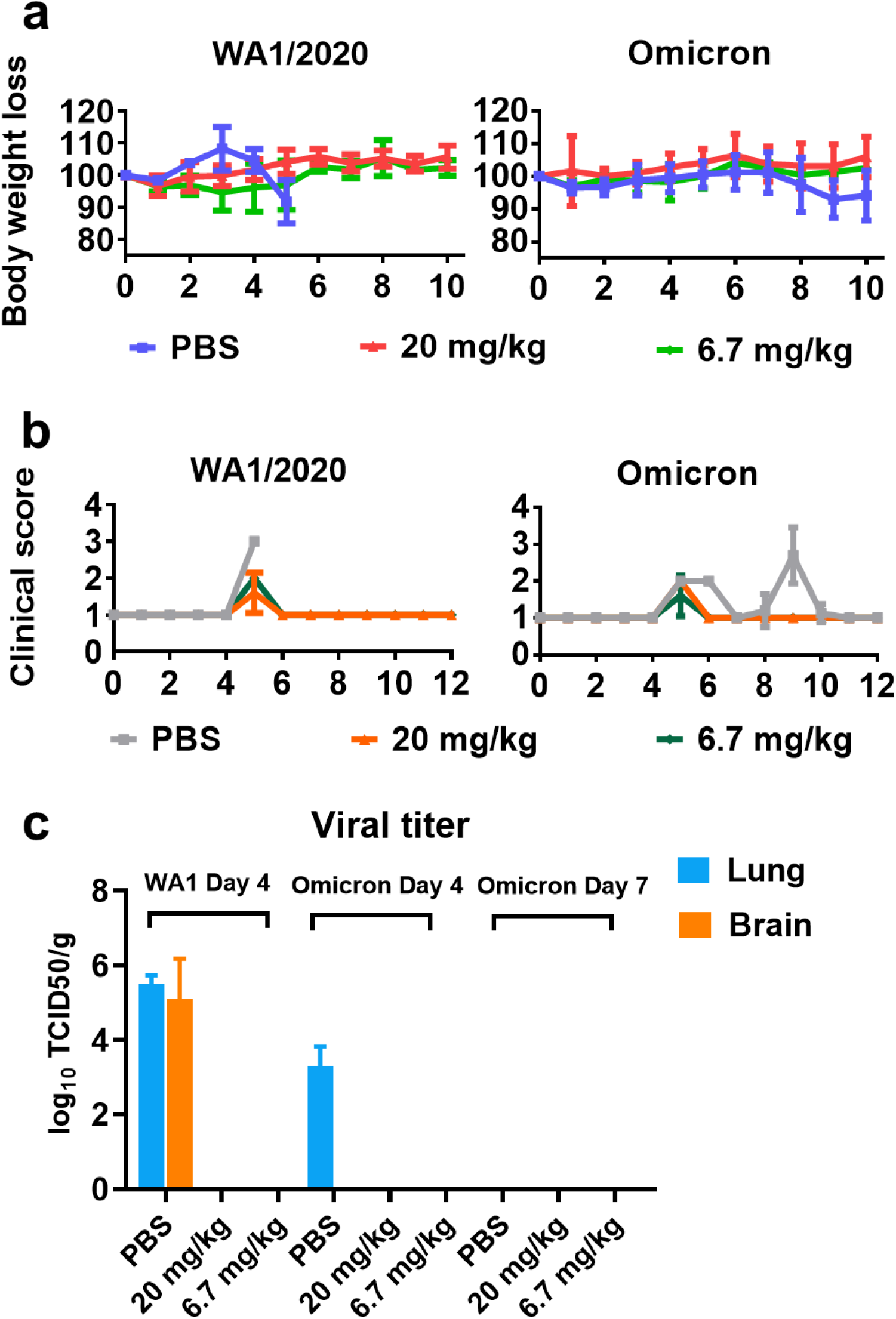
Therapeutic efficacy of antibody 8G3 agaisnt SARS-CoV-2 infection. a, Body weight change of mice. AC70 human ACE2 transgenic mice were challenged with 300 TCID_50_ of WA1/2020 or 2.3 × 10^4^ TCID_50_ of Omicron viruses and then received two doses of 8G3 treatment at 4 h and 48 h post infection. Clinical manifestations were observed at least once daily. b, Clinial score of mice. The clinical wellbeing of mice was assessed based on a 1-4 grading system. c, Viral titer in lung and brain tissues at day 4 and day 7. Data are shown as mean ± S.D.

## Discussion

There are more than 30 mutation sites on the entire S protein of SARS-CoV-2 Omicron, nearly half of which are distributed on the RBD region. These mutations together reshaped the S confirmation, increased the stability, enhanced its binding affinity with ACE2 receptor, as well as assisted viral evasion from targeted immunotherapy^13-16^. The mutation related changes greatly affects the transmissibility of the virus, the symptoms of infection, and host immune response^17-19^. In addition to general immune evasion from neutralizing antibodies, Omicron also broadly caused dramatic reduction in the protective efficacy of vaccines and sera^20-22^, which leads to a reduction in the options of responding to the SARS-CoV-2 pandemic and forces the generation of more passive prevention policies. Many neutralizing antibodies are disabled due to the large number of and widely scattered mutation sites in the N-terminal domain (NTD) and RBD region of the Omicron variant^23,24^. We investigated a memory B cell repertoire developed from early COVID-19 convalescents, and found that most of the WA1/2020 neutralizing antibodies had partially or completely lost their activities against Omicron. Antibody 8G3 neutralized Omicron more efficient than WA1/2020.

Several studies have reported the escape of EUA antibodies using pseudo-typed Omicron^6,7,25^. We compared the authentic Omicron neutralizing activities of several EUA antibodies targeting different epitope groups and confirmed that only Sotrovimab that has no epitope overlap with the RBD-ACE2 interface retained partial activity. Huge number of mutations in Omicron that reshape the surface of the S protein may shift the spatial positions of atoms and lead to changes in antibody binding, which brings more uncertainty to the prediction on antibody activity. Still and all, antibody 8G3 showed strong activity against Omicron in both in vitro neutralizing systems. We challenged transgenic mice with Omicron virus and found that the symptoms of infection of which were less severe than with WA1/2020 virus, and a proportion of mice recovered without antibody treatment. This might suggest a weakened virulence of the Omicron variant. Treatment with both 20 mg/kg and 6.7 mg/kg of antibody 8G3 quikly cleared the virus load with no clinical deterioration, demonstrating the potent therapeutic efficacy of 8G3.

The mutations of Omicron are mainly distributed in the front of the interface with ACE2 interaction (Fig. 2c). This could be used to explain why most of current EUA antibodies have generally lost neutralization. The development of a new class^26^ of neutralizing antibodies targeting conserved epitopes away from the ACE2 interface could be a viable approach to address Omicron escape^5^. The epitope of the antibody 8G3 is currently unknown. In view of its non-attenuated activity against Omicron, 8G3 might be a conserved epitope antibody that avoids the extensive mutation site distribution on Omicron. In summary, despite the universal desensitization of EUA antibodies caused by the emergence of Omicron, our results suggest that the use of antibodies with broad neutralization to address SARS-CoV-2 mutations remains a feasible strategy.

## Materials and methods

### Antibody preparation

The following procedure was used for all antibodies candidate’s generation. Plasmids containing the full-length of antibody heavy chain or light chain were amplified in *E. coli* DH5α and extracted by using PureLink™ HiPure Plasmid Miniprep Kit (Invitrogen). Antibodies were expressed using the ExpiCHO™ expression system (Thermo Fisher). The cell culture process was implemented following the manufacturer’s recommendations. The cell culture medium was collected 12 days post transfection and centrifuged at 300×g for 10 min to remove the cells. After further centrifugation at 7000×g for 30 min for removing the cell debris, the supernatant was subjected to MabSelect Sure (Cytiva) antibody affinity purification. In an AKTA avant protein separation and purification system (Cytiva), the MabSelect Sure column was equilibrated with 2 column volumes of 20 mM PB + 150 mM NaCl (pH 7.2). Then the medium samples were loaded and washed with 10 column volumes of 100 mM citric acid (pH 5.0). Antibodies were eluted with 5 column volumes of 100 mM citric acid (pH 3.0), followed by storage in a cocktailed buffer containing 10 mM Histidine-HCl, 9% trehalose, and 0.01% polysorbate 80.

Antibody biosimilars of Etesevimab, Casirivimab, Imdevimab, Sotrovimab and Bamlanivimab were purchased from ProteoGenix.

### 293T-ACE2 cells

Human ACE2 protein was stably expressed on the surface of HEK-293T cells (ATCC) to obtain the 293T-ACE2 cells. In this work, a three-plasmid lentivirus transfection system was used. The ACE2 gene was inserted into the pHIV-puro plasmid, and prepared together with the packaging plasmids psPAX2 and VSV-G using an Endo-free Plasmid Mini Kit (Omega). HEK-293T cells were seeded into a 10 cm dish and cultured at 37 °C under 5% CO_2_ condition. After the cells had grown to 70%-80% confluence, plasmids were co-transfected into cells using Lipofectamine 3000 reagent (Thermo Fisher). The cell medium was replaced with fresh DMEM medium (containing 10% FBS) 6 h later. After further culture for 48 h, the medium supernatant was collected and centrifuged at 3000×g for 10 min. The supernatant was transferred to a 15 mL centrifuge tube and mixed with the Lentivirus Concentration Reagent (Genomeditech) for overnight incubation. The next day, the mixture was centrifuged at 3220×g for 30 min, followed by resuspending the viral particles with 200 μL DMEM medium.

For infecting HEK-293T cells, 5 × 10^5^ cells were seeded into a 6-well plate and incubated overnight at 37 °C with 5% CO_2_. Then 10 μL of concentrated virus was added. 48 h later, cells were transferred to a 10 cm dish, and puromycin at final concentration of 10 μg/mL was added. Fluorescence activated cell sorting was used to further select the cells with high expression of ACE2. Using S1-mFc recombinant protein (Sino Biological) as the primary antibody and FITC-labeled goat anti-mouse IgG antibody (Jackson) as the secondary antibody, the cells in the top 1% of fluorescence intensity were obtained on a BD FACSJazz cell sorter (BD). The top 1% cells were expanded for pseudovirus neutralization experiments.

### Pseudovirus packaging and neutralization

The S sequences of SARS-CoV-2 (GenBank: QHD43416.1) or variants with intracellular 21 residue deletion were inserted into the pMD2G plasmid. A luciferase gene was constructed into plasmid pLVX-N1 as a reporter. Pseudoviruses were obtained as the lentivirus packaging method described above. For testing pseudovirus neutralizing activity by antibodies, 293T-ACE2 cells were seeded into a white 96-well plate (Corning) at a density of 1 × 10^4^ per well, and cultured overnight at 37 °C with 5% CO_2_. Serial 10-fold dilutions of antibodies from 200 μg/mL were mixed with equal volume of diluted pseudoviruses in DMEM medium (with 10% FBS). Medium containing the same amount of pseudovirus but without antibody was used as infection control. After incubation at 37 °C for 30 min, the medium of the 293T-ACE2 cells was replaced with 100 μL of antibody-pseudovirus mixture. All operations involving pseudoviruses were performed at a biosafety level 2 laboratory in Shanghai Jiao Tong university. After further culturing for 48 hours, 50 μL of ONE-Glo™ Luciferase Assay System substrate (Promega) was added to each well, and the fluorescence intensity was immediately measured in a microplate reader (TECAN). Data were analyzed using GraphPad Prism software, and the IC_50_ values were calculated using a four-parameter nonlinear regression function.

### Authentic virus neutralization

Infection with SARS-CoV-2 causes cell lysis especially in Vero E6 cells, which results in a cytopathic effect (CPE). Thus, the infection of the virus can be assessed by monitoring the CPE of the monolayer Vero E6 cells^27,28^. We employed a standard authentic virus-based micro-neutralization assay to assess the effect of neutralizing antibodies against SARS-CoV-2 infection. Vero E6 cells (ATCC) were seeded into a 96-well plate to reach approximately 85-90% confluence prior to infection. Antibodies were 3-fold serially diluted starting from a final concentration of 100 μg/mL and incubated with the cells for 1 h. SARS-CoV-2 WA1/2020-(US_WA-1/2020 isolate), Alpha-(B.1.1.7/UK, Strain: SARS-CoV-2/human/USA/CA_CDC_5574/2020), Beta-(B.1.351/SA, Strain: hCoV-19/USA/MD-HP01542/2021), Gamma-(P.1/Brazil, Strain: SARS-CoV-2/human/USA/MD-MDH-0841/2021), Delta-(B.1.617.2/Indian, Strain: GNL-751, a recently isolated strain from Galveston County, Texas) provided through World Reference Center for Emerging Viruses and Arboviruses (WRCEVA), chimeric between WA-1/S-Mu (SARS-CoV-2 Spike Col-B.1.621), infectious clone of Delta plus (icSARS-CoV-2 AY4+Y145H) and Omicron isolate (Strain: EHC_C19_2811c) were used in this study. Each strain was expanded in Vero E6 cells and resulting cell-free supernatants were sequenced, titrated, and stored at – 80 °C as working stocks. Only Passage #3 of SARS-CoV-2 (US_WA-1/2020) and Passage #1 of each variants were used for in vitro studies. Viruses in 100 particles per sample were added in duplicate. Medium with or without virus was used as positive and negative controls, respectively. The CPE in each well was observed under a microscope 4 days post infection. The IC_50_ values were calculated according to the following equation: IC_50_ = Antilog (D – C × (50 - B) / (A - B)). The A represents the inhibition rate when above 50%, B represents the inhibition rate when below 50%, C is log_10_ (dilution factor), D is log_10_ (sample concentration) when the inhibition is below 50%. All authentic SARS-CoV-2 studies were conducted under a Notification of Use (NOU) protocol approved by Institutional Biosafety Committee (IBC), at a biosafety level 3 laboratory in the Galveston National Laboratory, the University of Texas Medical Branch, Galveston, TX.

### Animal protection

Permissive AC70 human ACE2 transgenic mice (Taconic Biosciences) were intranasally challenged with 300 half tissue culture infectious dose (TCID_50_) of WA1/2020 virus (US_WA-1/2020) or 2.3 × 10^4^ TCID_50_ of Omicron virus (Strain: EHC_C19_2811c). After that, mice received two doses of antibody 8G3 (20 mg/kg or 6.7 mg/kg) or equal volume of PBS by tail vein injection at 4 h and 48 h post infection. On day 4, three mice in each group were sacrificed for viral load and histopathology examination. On day 7, another three mice in the Omicron-infected groups were euthanized for the second viral load and histopathology examination. Clinical wellbeing and body weight of the remaining five mice in each group were observed and recorded at least daily, until the designed endpoint at day 14. The performances of mice were scored based on a 1 to 4 grading system, where score 1 indicates healthy, score 2 indicates with ruffled fur and lethargic, score 3 indicates with additional clinical sign such as hunched posture, orbital tightening, increased respiratory rate, and/or > 15% weight loss, and score 4 indicates dyspnea and/or cyanosis, reluctance to move when stimulated, or ≥ 20% weight loss that need immediate euthanasia. Animal studies were carried out at Galveston National Laboratory at University of Texas Medical Branch at Galveston, Texas, based on a protocol approved by the Institutional Animal Care and Use Committee at UTMB at Galveston.

